# Inhibition of dopamine neurons prevents incentive value encoding of a reward cue: With revelations from deep phenotyping

**DOI:** 10.1101/2023.05.03.539324

**Authors:** Amanda G. Iglesias, Alvin S. Chiu, Jason Wong, Paolo Campus, Fei Li, Zitong (Nemo) Liu, Shiv A. Patel, Karl Deisseroth, Huda Akil, Christian R. Burgess, Shelly B. Flagel

## Abstract

The survival of an organism is dependent on their ability to respond to cues in the environment. Such cues can attain control over behavior as a function of the value ascribed to them. Some individuals have an inherent tendency to attribute reward-paired cues with incentive motivational value, or incentive salience. For these individuals, termed sign-trackers, a discrete cue that precedes reward delivery becomes attractive and desirable in its own right. Prior work suggests that the behavior of sign-trackers is dopamine-dependent, and cue-elicited dopamine in the nucleus accumbens is believed to encode the incentive value of reward cues. Here we exploited the temporal resolution of optogenetics to determine whether selective inhibition of ventral tegmental area (VTA) dopamine neurons during cue presentation attenuates the propensity to sign-track. Using male tyrosine hydroxylase (TH)-Cre Long Evans rats it was found that, under baseline conditions, ∼84% of TH-Cre rats tend to sign-track. Laser-induced inhibition of VTA dopamine neurons during cue presentation prevented the development of sign-tracking behavior, without affecting goal-tracking behavior. When laser inhibition was terminated, these same rats developed a sign-tracking response. Video analysis using DeepLabCut revealed that, relative to rats that received laser inhibition, rats in the control group spent more time near the location of the reward cue even when it was not present and were more likely to orient towards and approach the cue during its presentation. These findings demonstrate that cue-elicited dopamine release is critical for the attribution of incentive salience to reward cues.

**Significance Statement:** Activity of dopamine neurons in the ventral tegmental area (VTA) during cue presentation is necessary for the development of a sign-tracking, but not a goal-tracking, conditioned response in a Pavlovian task. We capitalized on the temporal precision of optogenetics to pair cue presentation with inhibition of VTA dopamine neurons. A detailed behavioral analysis with DeepLabCut revealed that cue-directed behaviors do not emerge without VTA dopamine. Importantly, however, when optogenetic inhibition is lifted, cue-directed behaviors increase, and a sign-tracking response develops. These findings confirm the necessity of VTA dopamine during cue presentation to encode the incentive value of reward cues.

## Introduction

Associative learning strategies are utilized daily by humans and animals alike to make situational decisions. Such strategies often rely on cues, or stimuli, in the environment to guide behavior and can directly impact the survival of an organism. In rodents, individual differences in cue-motivated behaviors can be captured using a Pavlovian conditioned approach (PavCa) paradigm, wherein presentation of a discrete cue (conditioned stimulus, CS) is followed by delivery of a food reward (unconditioned stimulus, US) (Flagel et al., 2009). Following PavCa training, two distinct phenotypes emerge – goal-trackers (GT) and sign-trackers (ST) (Boakes, 1977; Hearst, 1974; Robinson & Flagel, 2009). While both GTs and STs attribute predictive value to the reward cue, STs also attribute incentive value to the cue. The attribution of incentive motivational value, or incentive salience, transforms the cue itself into an attractive and desirable stimulus (Berridge & Robinson, 2003). For STs, both food- and drug-associated cues gain appreciable incentive value and thereby the ability to elicit maladaptive behaviors (Saunders & Robinson, 2010, 2011; Yager et al., 2015; Yager & Robinson, 2013). The ST/GT model, therefore, can be harnessed to elucidate the neurobiological mechanisms that encode the predictive versus incentive value of reward cues. Further, this model can help us better understand the neural processes that contribute to shared symptomatology between psychiatric disorders, as an increased propensity to attribute incentive salience to reward cues (i.e. to sign-track) has been associated with externalizing behaviors and deficits in executive control in both rodents and humans (Colaizzi et al., 2023; Flagel et al., 2010; Phillips & Sarter, 2020).

Dopamine has been implicated in a number of psychiatric disorders, predominantly via its role in learning, attention, and motivation (Grace, 2016; Howes & Kapur, 2009; Nestler & Carlezon, 2006; Volkow et al., 2017). However, the precise role of dopamine remains a subject of debate, especially as it pertains to reward processing and learning about stimuli in the environment (Berke, 2018; Berridge, 2007; Lerner et al., 2021; Schultz et al., 2017). While dopamine has long been considered a prediction error signal (Schultz et al., 1997), used to update the predictive value of reward-cues during associative learning, deficiencies in this theory have been recognized (e.g., Jeong et al., 2022; Kutlu et al., 2021; Saunders et al., 2018; Sharpe et al., 2020). Of particular relevance, the sign-tracker/goal-tracker model previously revealed that intact dopamine signaling is necessary for the attribution of incentive salience to reward cues, or what we refer to here as Pavlovian “incentive learning”, and not the encoding of predictive value alone, or “predictive learning” (Flagel et al., 2011; Saunders & Robinson, 2012; Yager et al., 2015). Both systemic and local (nucleus accumbens core) blockade of dopamine receptors prevents the acquisition and expression of a sign-tracking conditioned response with no effect on goal-tracking (Flagel et al., 2011; Saunders & Robinson, 2012). Additionally, using fast-scan cyclic voltammetry, it was shown that dopamine-encoded “prediction error” signals are present in the nucleus accumbens core of sign-trackers, but not goal-trackers (Flagel et al., 2011). Together, these findings led to the conclusion that dopamine encodes the incentive value of Pavlovian reward cues. However, it was not clear from these studies whether dopamine activity precisely at the time of cue presentation is necessary for incentive value encoding and the acquisition of sign-tracking behavior. To address this question, we exploited the temporal resolution of optogenetics. Specifically, we utilized tyrosine hydroxylase (TH)-Cre rats to selectively inhibit dopamine neurons in the ventral tegmental area (VTA) during cue presentation early in Pavlovian training. We found that male TH-Cre Long Evans rats have an inherent tendency to sign-track, and that optogenetic inhibition of VTA dopamine neurons during cue presentation blocks this tendency, without affecting goal-tracking behavior. An in-depth analysis of behavior using DeepLabCut (Mathis et al., 2018) revealed that the effects of this manipulation were time-locked and specific to cue-elicited incentive motivation.

## Materials and Methods

### General Methods

#### Subjects

One hundred twenty-eight male Long Evans rats were received from a breeding colony maintained by Dr. Huda Akil’s laboratory (Michigan Neuroscience Institute, University of Michigan, Ann Arbor, MI) (Figure 1a). Only male rats were used for these studies, which are a direct follow-up to prior studies that had been conducted with male rats (Flagel et al., 2011; Saunders & Robinson, 2012). The breeding colony originated in 2013 with two TH-Cre male Long Evans rats from the Deisseroth laboratory (Stanford University, Stanford, CA). The colony has since been maintained by breeding TH-Cre Long Evans male rats with wild type (WT) Long Evans female rats. Rats were bred and weaned in either the Biological Science Research Building or the Molecular and Behavioral Neuroscience Institute building (University of Michigan, Ann Arbor, MI). Rats were transferred to the Flagel laboratory in the Molecular and Behavioral Neuroscience Institute building around postnatal day (PND) 46. They were housed under a 12-hour light-dark cycle (lights on at 6 or 7 AM depending on daylight saving time) with climate-controlled conditions (22 ± 2°C). Rats had *ad-libitum* access to food and water throughout the study. They were pair- or triple-housed prior to surgery, and single-housed following surgery and for the duration of the experiment. Rats were acclimated to the housing room for a minimum of one week before experimenter handling began. Behavioral testing took place during the light-phase between 10 AM and 4 PM. All procedures followed The Guide for the Care and Use of Laboratory Rats: Eighth Edition (2011, National Academy of Sciences) and were approved by the University of Michigan Institutional Animal Care and Use Committee.

**Figure 1.**
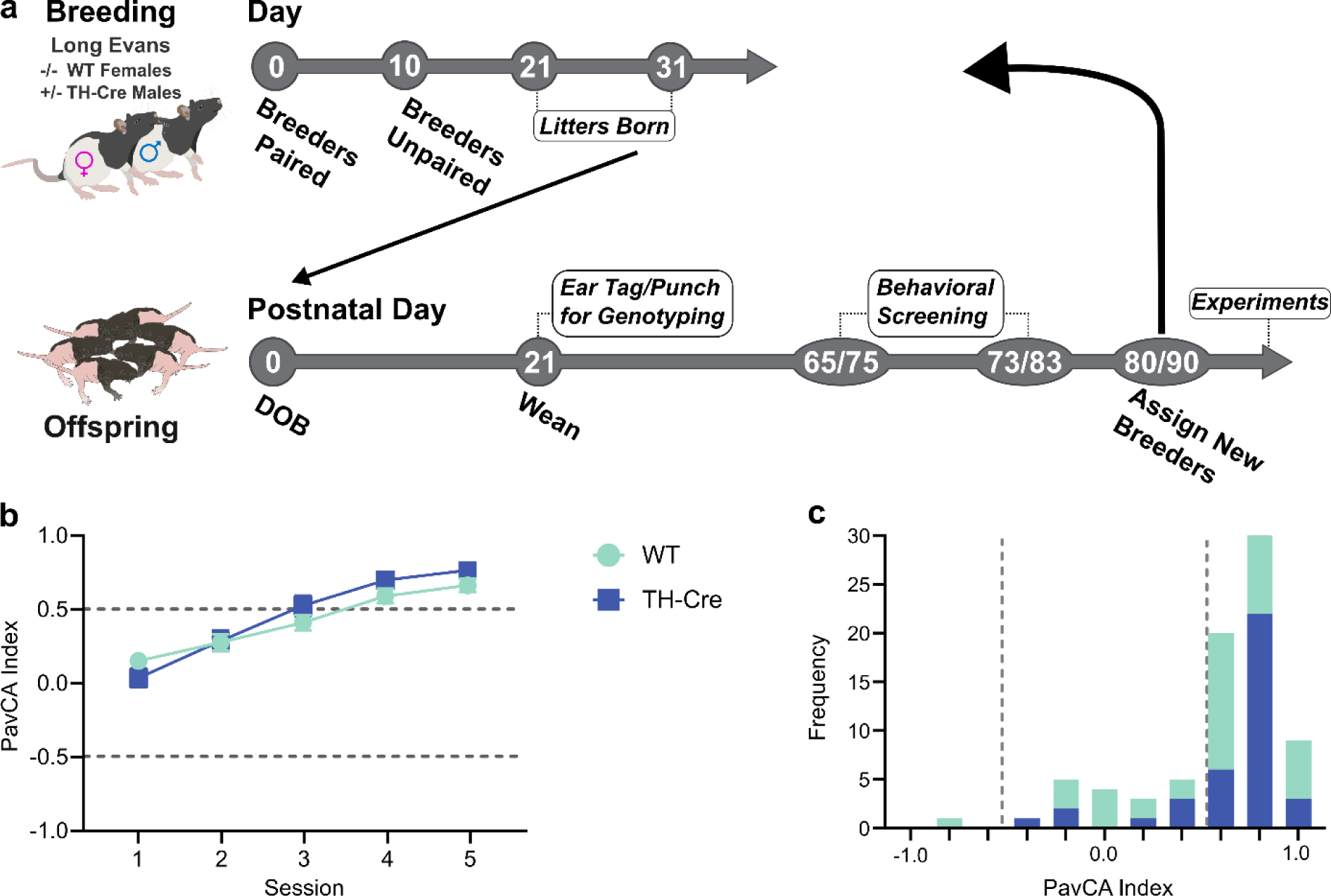
The population of Long Evans male rats are skewed toward sign-trackers. (a) Breeding schematic and timeline depicting maintenance of the Long Evans TH-Cre breeding colony. (b) Pavlovian conditioned approach (PavCA) Index shown as mean ± SEM across 5 training sessions for WT rats (n = 57) and naïve TH-Cre rats (n = 38). (c) Frequency histogram illustrating the number of rats exhibiting a mean PavCA Index (averaged from sessions 4 and 5) between −1.0 and +1.0 for each of the groups depicted in panel b. 84% of the population (n = 95) was skewed toward sign-trackers (≥ 0.5, n = 46 WT and n = 34 TH-Cre), 14% of the population were intermediate responders (0.4 ≤ −0.4; n = 10 WT and n = 3 TH-Cre, cutoff 0.4 ≤ −0.4), and 2% goal-trackers (≤ −0.5, n = 1 WT and n = 1 TH-Cre).

To determine the inherent behavioral phenotypes of the transgenic rats (Figure 1), 95 male rats were used. The TH-Cre (n = 38) rats were from 2 generations and 7 litters, and the WT (n = 57) rats were from 2 generations and 15 litters. To determine the effects of optogenetic inhibition of dopamine neurons during Pavlovian cue-reward learning (Figure 2), 33 male TH-Cre rats from 3 generations and 11 litters were used. Some rats were excluded for not consuming pellets during pretraining (n = 4), poor virus expression/probe placement (n = 8), or head caps coming off prematurely (n = 4). Due to technical issues, session 4 data was lost for one rat in the halorhodopsin group and session 5 data was lost for one rat in the control group. Data from these two rats are included in the analyses for other sessions. In total, 17 out of 33 rats are included in the behavioral analyses assessing the effects of optogenetic inhibition, with 10 in the halorhodopsin group and 7 in the control group. For DeepLabCut analyses, three of these rats were excluded due to technical issues with video capturing, resulting in 8 in the halorhodopsin group and 6 in the control group.

**Figure 2.**
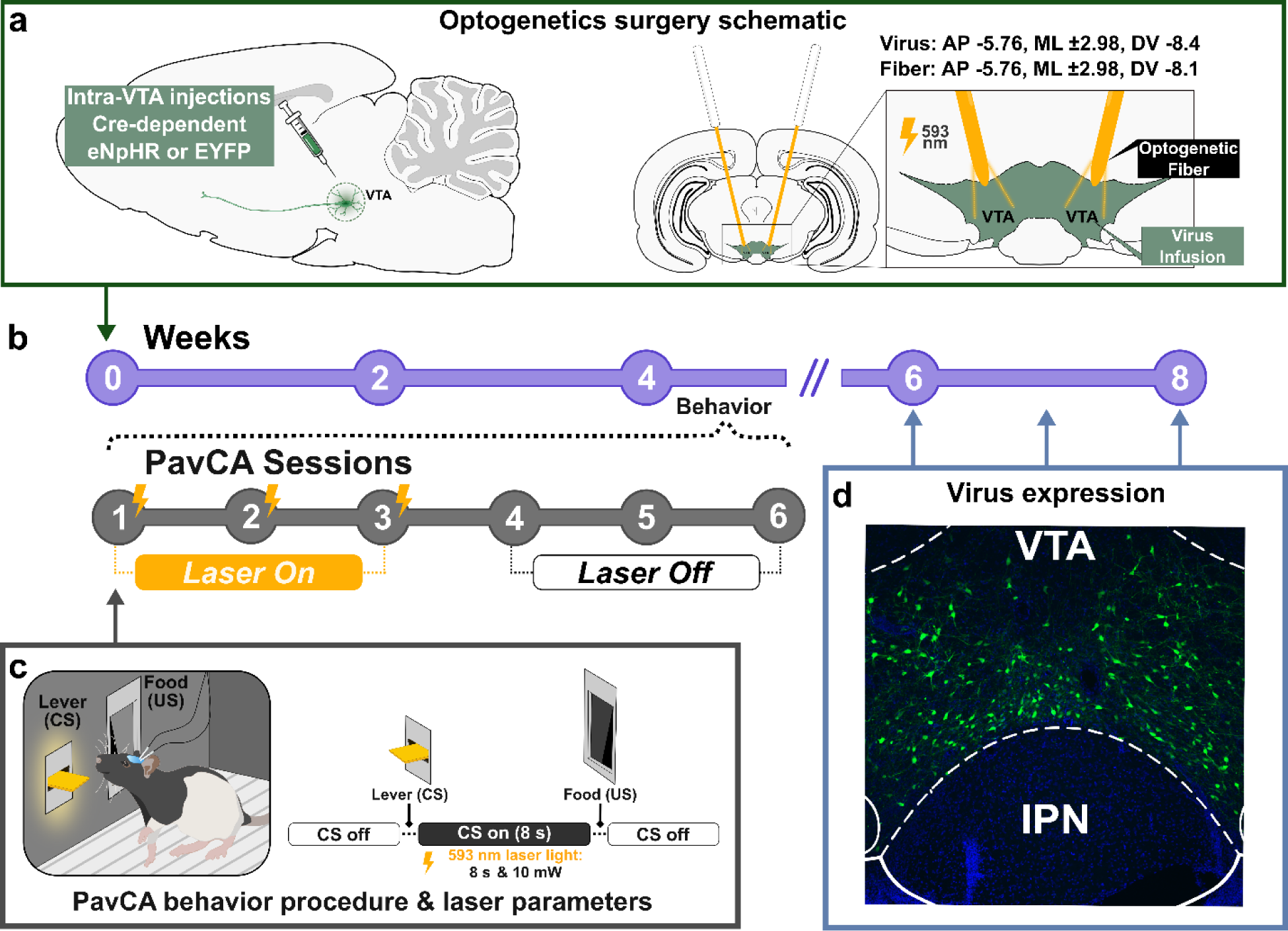
Experimental timeline. (a) Schematic illustrating optogenetic viral infusion and fiber placement. 1.0 µl of either eNpHR (halorhodopsin) or EYFP (control) virus was infused and the optogenetic fiber was placed bilaterally in the ventral tegmental area (VTA). (b) Week of experimental timeline represented in purple. Week 4 of the experiment is expanded below in gray to show Pavlovian conditioned approach (PavCA) sessions. (c) Schematic of behavioral paradigm in which lever-CS presentation (8.2 sec) was paired with laser inhibition of dopamine neurons during PavCA sessions 1-3. The laser remained off for the last 3 PavCA sessions. (d) Representative image of virus expression in the VTA.

#### Surgery

All rats had anesthetic induction of 5% isoflurane (Vet One, Boise, ID) delivered via an induction chamber. Following induction, artificial tears (Akorn, Gurnee, IL) were placed over the eyes and an area of the rat’s scalp was shaved between the posterior edge of the eyes and the ears. Rats were placed into the stereotaxic frame (David Kopf instruments, Tujunga, CA or Stoelting, Wood Dale, IL), given an injection of carprofen (5 mg/kg, subcutaneous (s.c.)) for analgesia, and the shaved scalp was cleaned with betadine (Veterinary Betadine, Stamford, CT) and alcohol. An incision on the scalp exposed the skull. A drill (Foredom, Bethel, CT) was secured to the stereotax and used for leveling the skull such that the tip of the drill bit was zeroed at bregma (Paxinos & Watson, 2007). Rats were levelled in the stereotaxic frame by adjusting the ear bars. The drill bit was moved ± 2 mm ML (medial/lateral) from bregma and lowered to the skull. The DV (dorsal/ventral) coordinate for each site was recorded. If the difference between the sites was greater than ± 0.05 mm DV the animal was adjusted. Following ML adjustment, the drill bit was moved to lambda (Paxinos & Watson, 2007) and the angle of the nose was adjusted until bregma and lambda were within ± 0.05 mm DV.

#### Pavlovian conditioned approach (PavCA)

Prior to assessing Pavlovian conditioned approach (PavCA) behavior, rats were handled by the experimenters. Those used for screening the colony were handled for 2 days and those used for the optogenetics experiment were handled for 4 days. As described below, a single pretraining session preceded PavCA sessions. For the two days prior to pre-training rats received ∼25 banana-flavored grain pellets (each pellet 45-mg; Bio-Serv, Flemington, NJ, USA) in their home cage to acquaint them with the reward.

PavCA testing occurred inside Med Associates chambers (St. Albans, VT, USA; 24.1 × 21 x 30.5 cm) located in sound-attenuating boxes equipped with a fan to reduce background noise (Figure 2c). The chambers contained a magazine that was connected to a pellet dispenser and placed in the center of one wall 3 cm above the chamber floor. An illuminated retractable lever (i.e. lever-cue) was located either to the left or right of the magazine (Med Associates, ENV-200R2M-6.0), 6 cm above the chamber floor. The magazines used were taller (2 x 6 in) than the standard (2 x 2 in) to allow rats to access food pellets without interference from their headcap and tethers. A white house light was placed at the top of the chamber on the wall opposite to the magazine and lever-cue and remained on for the duration of the session. Magazine entries were recorded by a break of the photo beam inside the cup. Lever-cue contacts were recorded following a minimum force of 10 g.

Prior to each session the rats were transferred to the testing room in their home cage. They were left in the room for a minimum of 30 min to allow them to acclimate. Rats were initially placed into the Med Associates chambers for a single pretraining session. At the start of the pretraining session, the food magazine was baited with two food pellets to direct the rats’ attention to the location of reward delivery. During pretraining the lever-cue remained retracted, and rats received a food pellet in the food magazine on a variable 30 s (range 0-60 s) schedule. There was a total of 25 trials and the pretraining session lasted approximately 12.5 min wherein head entries were recorded, and food pellet consumption confirmed. Following pretraining rats had a single session of PavCA each day for either 5 (colony characterization) or 6 (optogenetic inhibition) consecutive days. The start of each PavCA session began with a 5-min waiting period followed by the house light turning on which signified the start of the session. As previously described (Campus et al., 2019; Meyer et al., 2012), during PavCA, the illuminated lever-cue (conditioned stimulus, CS) entered the chamber for 8 s and upon retraction a food pellet (unconditioned stimulus, US) was immediately delivered into the adjacent food magazine. PavCA sessions consisted of 25 lever-cue (CS)/ food-US trials on a variable 90-s schedule (range 30–150 s). Each session lasted approximately 40 min. It was confirmed that all food pellets had been consumed following each session.

Med Associates software recorded the following behaviors during PavCA sessions: 1) number of food magazine contacts made during the 8-s lever-cue presentation, (2) latency to contact the food magazine during lever-cue presentation, (3) number of lever-cue contacts, (4) latency to lever-cue contact, and (5) the number of food magazine entries made during the inter-trial interval (i.e., food magazine contacts made in between lever-cue presentations). Contact and latency data were used to calculate the PavCA Index to characterize rats into their behavioral phenotypes, as previously described (Meyer et al., 2012). The PavCA Index is a composite measure calculated using the following formula: [Probability Difference Score + Response Bias Score + (−Latency Difference Score)/3]. PavCA Index scores range from −1 to 1, with a score of −1 representing individuals with a conditioned response (CR) focused solely on the food magazine during lever-cue presentation (i.e., a “pure” goal-tracker, GT) and a score of +1 representing individuals with a CR focused solely on the lever-cue upon its presentation (i.e., a “pure” sign-tracker, ST) (Figure 1b). For colony characterization, the PavCA Index from sessions 4 and 5 were averaged to assess the frequency distribution of sign-trackers and goal-trackers (Figure 1c).

### Behavioral characterization of the Long Evans colony

#### Subjects

A subset (n = 95) of male Long Evans and TH-Cre rats underwent “baseline” characterization of PavCA behavior (Figure 1). Naïve WT rats (n = 26) and WT rats following sham surgery (n = 31) were included in this analysis. Sham surgery consisted of levelling rats in the stereotaxic frame and drilling two holes directly above the VTA (bilaterally from bregma, AP −5.76; ML ± 2.98). A 1 µL Hamilton Neuros Syringe was lowered into the holes (from bregma, DV −8.4) and pulled up after 10 min. The surgical site was closed with clips (Stoelting, Wood Dale, IL) which were removed 7-10 days following surgery. After the clips were removed experimenter handling and behavioral testing procedures began. TH-Cre rats without prior experimental manipulation (n = 38) were screened for “baseline” PavCA behavior and subsequently used for pilot studies.

#### Statistical analyses

To compare differences in the PavCA Index among the colony (Figure 1b), a linear mixed-effects model (LMM) with restricted maximum likelihood estimation was used. This analysis applies multiple covariance structures to the data set and the structure with the lowest Akaike information criterion (AIC) was selected as best fit (Duricki et al., 2016; Verbeke, 1997). LMM was conducted to compare genotypes and groups across sessions 1-5. Session was used as the repeated variable and genotype/group as the between-subjects variable. For one analysis the WT group was split into rats that were Naïve (WT Naïve) and those that received Sham surgery (WT Sham), and for another analysis, the WT groups were collapsed and compared to TH-Cre rats. To assess the differences in the relative proportions of phenotypes between the genotypes in the Long Evans colony, a Fisher’s Exact Test was conducted on the PavCA Index (Figure 1c). For all analyses, statistical significance was set at *p* < 0.05, and Bonferroni *post hoc* comparisons were made when significant main effects or interactions were detected.

### Optogenetic inhibition of the VTA

#### Viral vectors

A Cre-dependent inhibitory optogenetic construct halorhodopsin (eNpHR, AAV5-Ef1a-DIO eNpHR 3.0-EYFP at titer ≥ 1×10¹³ vg/mL, Addgene plasmid # 26966) or an empty vector (control, AAV5-Ef1a-DIO EYFP at titer ≥ 1×10¹³ vg/mL, Addgene plasmid # 27056) were utilized. Both plasmids were obtained from Addgene as gifts from Dr. Karl Deisseroth.

#### Virus and optogenetics probe implant surgery

Rats were around PND 91 at the time of surgery. Two holes were drilled directly above the VTA (bilaterally from bregma, AP −5.76; ML ± 2.98). Four additional holes were drilled ± 2 mm ML from bregma - two were directly behind the coronal suture and two were directly behind the lambdoid suture. 2.4 mm stainless-steel screws (Plastics One, Roanoke County, VA) were secured into the four holes. A Hamilton Syringe (5 µL Model 85 RN, Small Removable Needle, 26s gauge, 2 in, point style 2) was placed into a pump (Harvard Apparatus Pump 11 Elite, Holliston, MA) and then connected to P50 tubing and a guide cannula (Plastics One, Roanoke County, VA) that screwed onto a 10 mm injector (Plastics One, Roanoke County, VA). A Cre-dependent inhibitory optogenetic construct (halorhodopsin, eNpHR) or Cre-dependent control virus (EYFP) was bilaterally injected into the VTA at a 10° angle (from bregma, AP −5.76; ML ± 2.98; DV −8.4) at a rate of 100 nL per min over a 10-min period (1 µL total) (Figure 2a). The injector remained in place for an additional 10 min. After diffusing, fiber optic implants were inserted 0.3 mm above the injection site at a 10° angle (from bregma, AP −5.76; ML ± 2.98; DV −8.1, Figure 2a). Fiber optic implants were made in house and consisted of 200 µm-diameter optic fibers (Thor Labs, Newton, NJ) inserted into 10.5-mm-long ferrules (Thor Labs, Newton, NJ). Only fibers above 85% laser emittance were used for surgery. The fiber optic implants were secured with acrylic cement (Bosworth New Truliner, Keystone Industries, Gibbstown, NJ). The plastic screw from a guide cannula (Plastics One, Roanoke County, Virginia) was placed at the most anterior portion of the headcap as the acrylic cement was drying (later used for securing the rats headcap during behavior). A 3–4-week period for virus incubation followed surgery (Figure 2d).

#### PavCA sessions and laser parameters

Rats were handled by the experimenters for 4 days prior to the first session of behavior. The final two days of handling the rats received ∼25 banana-flavored grain pellets (each pellet 45 mg; Bio-Serv, Flemington, NJ, USA) in their home cage. Prior to testing the acrylic headcap was covered in black pet safe nail polish (Warren London Pawdicure Dog Nail Polish Pen) to occlude the laser light from illuminating the behavioral chamber. Prior to each PavCA session, rats were secured to the optogenetic probes. While being held by the experimenter the plastic screw at the front of the headcap was connected to a reinforced cannula spring (Plastics One, Roanoke, VA) and the optogenetic probes cleaned with ethanol and bilaterally connected to individual optogenetic cables that were secured with a ceramic mating sleeve (Thor Labs, Newton, NJ).

Rats had 1 pretraining session followed by 6 PavCA sessions. For the first 3 PavCA sessions (trials 1-75), rats received photoinhibition of the VTA continuously during the 8 s lever-cue presentation via a 593.5 nm Yellow DPSS Laser (Shanghai Laser & Optics Century CO., Ltd., Shanghai, China) (Figure 2b-d). Parameters known to be effective for inhibiting dopamine neurons were used (Gradinaru et al., 2010; McCutcheon et al., 2014). Laser power was calibrated to ∼10 mW/mm^2^ from the tip of the optogenetic cables before each session. Cables were also tested after each session to ensure laser power was consistent throughout. For sessions 4-6 (trials 76-150), rats were connected to the reinforced cannula spring and optogenetic cables as described above, but the laser was turned off (i.e., no photoinhibition occurred on sessions 4-6). Each rat had 3 PavCA sessions of “laser on” followed by 3 PavCA sessions of “laser off”, across 6 consecutive days.

#### PavCA orienting response

In addition to the behavioral metrics described above that were recorded with Med Associates software, behavioral outcome measures were also obtained via Experimenter observation of videos from session 3 of PavCA. Specifically, an Experimenter (A.G.I.) assessed whether a rat oriented towards the lever-cue or food magazine for each trial. An orienting response was defined as a head movement directed towards the lever-cue or food magazine at any point during the 8.2 sec lever-cue presentation. The probability of approaching either the lever-cue or food magazine was then calculated as the number of trials with an orienting response (to either the lever-cue or food magazine)/25. In addition, the percentage of trials on which an orienting response was directed towards the lever-cue, food cup, or both was determined.

#### Perfusion and tissue processing

Rats were perfused within 5 days following the experiment. Rats were first anesthetized with ketamine (90 mg/kg, intraperitoneal (i.p.)) and xylazine (10 mg/kg, i.p.) and then transcardially perfused with 0.9% saline and 4% formaldehyde (pH = 7.4). Following brain extraction, the tissue was post-fixed in 4% formaldehyde for 24 hours at 4°C and then placed in 30% sucrose at 4°C (sucrose in 0.1M PBS, pH = 7.4) for 3 days. The brains were frozen using dry ice and coated in a Tissue-Plus Optimal Cutting Temperature compound (Fisher HealthCare, Houston, TX). Coronal brain slices were taken at 40 µm using a cryostat at −20°C (Leica Biosystems Inc, Buffalo Grove, IL). The whole brain was collected, and slices were placed into well plates containing cryoprotectant and then stored at −20°C until further processing. Slices with the VTA were isolated, mounted onto SuperFrost Plus microscope slides (Fisher Scientific), and cover slipped with DAPI as a counterstain (diluted 1:5000 in 90% glycerol). Images were captured using a Zeiss AxioImager M2 motorized fluorescent microscope (Carl Zeiss, Sweden). Fluorescent images of endogenous virus expression and optogenetic probe placement were evaluated by two experimenters blind to the experimental groups (Figure 2d, representative image). Virus expression was evaluated based on distinct localized cell body expression of EYFP (virus tag) within the VTA (e.g., Figure 2d) and probe placements were confirmed if they were visualized bilaterally within the VTA (Figure 3).

**Figure 3.**
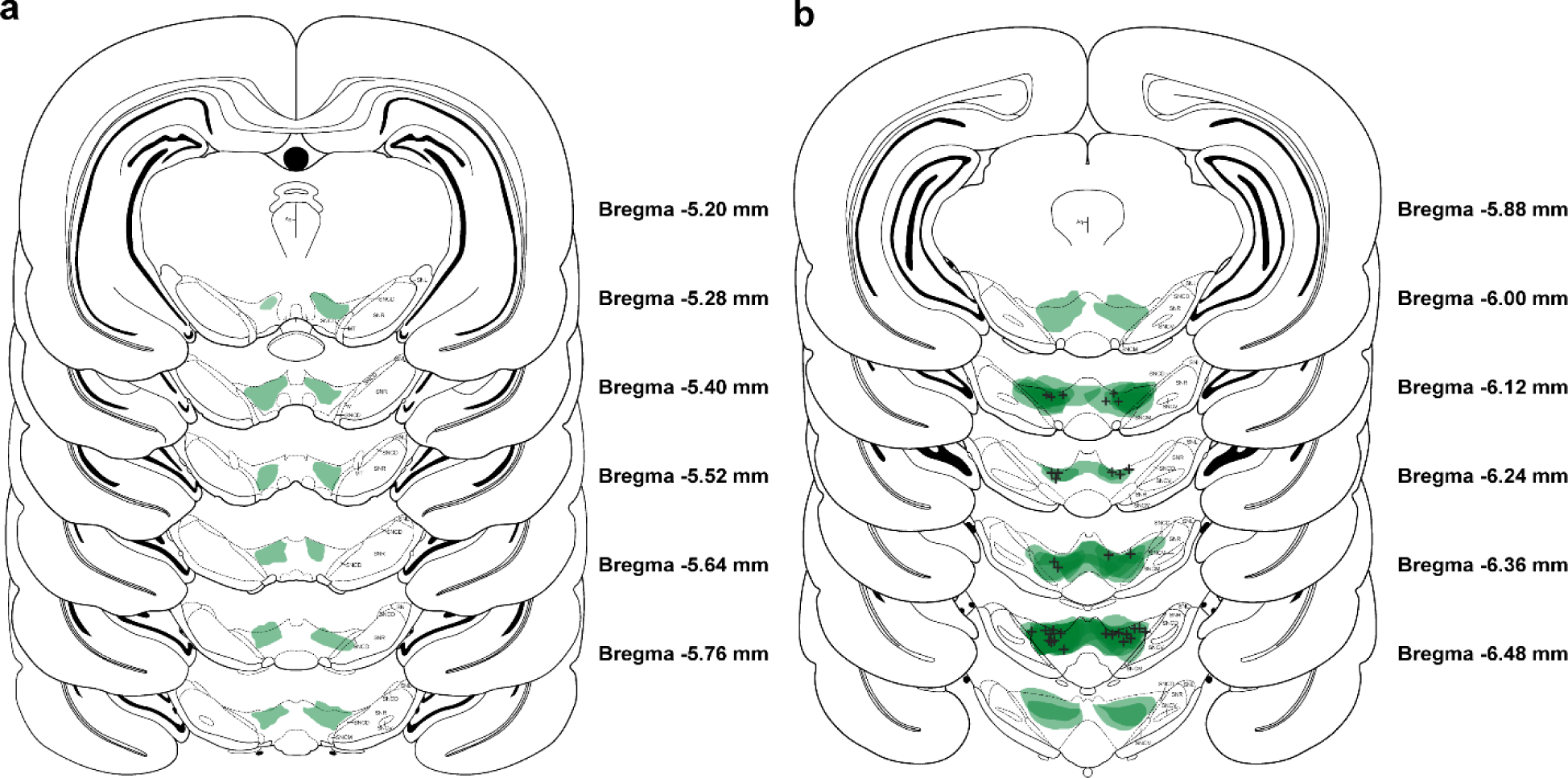
Virus expression and optogenetic probe placement in VTA. Coronal atlas images, (a) anterior to (b) posterior, relative to Bregma −5.20 to −6.48 are depicted with virus expression spread (in green) and probe placement markers (black + sign) for both experimental groups (halorhodopsin and EYFP). The density of the virus expression is reflected by the different hues of green, with darker color reflecting greater density. The densest virus expression and successful probe placements were between Bregma −6.12 to −6.36.

#### DeepLabCut Video Analysis

Videos from session 3 of PavCA were analyzed using DeepLabCut (DLC) and custom MATLAB (R2021b) scripts by experimenters (A.S.C and N.L.) who were blind to experimental groups. Raw videos were processed in Adobe Premiere Pro to increase contrast and enhance brightness within the behavioral chambers. Videos in this dataset on average contained 35,000 frames of which 75 frames were extracted for training. Training videos were from session 1. Labeling of all videos for training was completed by two experimenters (A.S.C and N.L.). Each video was manually labeled with specific markers (Figure 6a) of relevance to the planned analyses, including: tether, nose, left-ear, right-ear, left-shoulder, right-shoulder, and tail base on the rats. The food magazine and lever-cue location in the chamber were also labeled. Training was conducted via DeepLabCut 2.1.10.4 downloaded from GitHub (https://github.com/DeepLabCut/DeepLabCut) and installed onto University of Michigan, Great Lakes Computing Cluster. Following training of the network, locations of each marker for images analyzed on separate videos were extracted with a p-cutoff-parameter of 0.8. Videos for analysis that failed to meet criteria (>10 outlier frames) were reanalyzed following relabeling of outlier frames and retraining of the network. Three videos were excluded from analyses due to camera angle (one control rat and one halorhodopsin rat) or an occluded camera (one halorhodopsin rat).

Data extracted from session 3 videos were post-processed in MATLAB through linear interpolation and alignment with Med Associates Data. Linear interpolation was applied in addition to DLC filtering to further smooth pose estimation coordinates for frames during which body markers were not visible due to lighting or occlusion. Timepoints of each lever-cue presentation per trial were extracted from analyzed videos to validate alignment to lever-cue presentation based on Med Associates data. The following behaviors were extracted and analyzed in MATLAB: time in zone (food magazine or lever-cue), head orientation to food magazine or lever-cue, approach behavior (approach bouts) towards food magazine or lever-cue, locomotion, and latency to approach the food magazine. The length of the time bins for illustration and analyses were different across outcome measures to better capture behaviors that were time-locked to cue-presentation or retraction, and/or those that occurred during the intertrial intervals. Further, the reference body part differs for some of the measures as described below. For time in zone, body position was tracked by following the rats’ headcap tether throughout the chamber. The tether was chosen as the reference point due to the robustness of this marker tracking, compared to others, prior to interpolation. Each heatmap (time in zone, Figure 6b-e) represents the average location of rats across all trials on session 3, integrated over 123 video frames (8.2-s time bins) immediately before lever-cue presentation (Figure 6b), during lever-cue presentation (Figure 6c), after lever-cue presentation (Figure 6d), and during the intertrial interval (ITI, Figure 6e). A 2D Gaussian smoothing kernel with a standard deviation of 5 was applied to smooth the heatmap images (Figure 6b-e). Head orientation (Figure 7a) was tracked by angle changes between two vectors: 1) from the points between the rats’ nose to headcap tether and 2) from the rats’ headcap tether to the food magazine. For head orientation, the center of the food magazine was the reference point, and the videos were assessed across 4-s time bins during different phases of the session: 4-s before lever-cue presentation (Figure 7b), the first and last 4-s of lever-cue presentation (Figure 7c), the 4-s immediately after pellet delivery (Figure 7d), and 4 s during the ITI (Figure 7d). For approach behavior (Figure 8), body position was tracked by following the rats’ nose, the reference body part for this metric, throughout the chamber. An approach bout was counted when the rats nose remained within 75 pixels (1 cm) of the food magazine or lever-cue during lever-cue presentation (Figure 8a) and after lever-cue presentation (Figure 8b) for more than 15 frames (1 s). The bout ended when the interbout interval exceeded 15 frames (1 s). Locomotion (Figure 9) reflects the distance travelled based on tracking the rats’ center of mass, the point between the rat’s right and left shoulders (e.g., Figure 6a). Using the above indices, non-specific locomotion was averaged across the session (Figure 9a), lever-cue-elicited locomotion was captured during the first 2 s of lever-cue presentation (Figure 9b), and latency to approach the food magazine (Figure 9c) was assessed following lever-cue retraction.

#### Code Accessibility

DLC settings, desktop parameters, and code for performing post-processing reconstructions and analysis are made publicly available on GitHub: https://github.com/alvchiu/THCRE-dlc-.

#### Statistical analyses

Statistical analyses were conducted with the Statistical Package for the Social Sciences (SPSS) program version 27.0 (IBM, Armonk, NY, USA). To assess PavCA behavioral outcome measures across sessions, a linear mixed-effects model (LMM) with restricted maximum likelihood estimation was used to account for repeated measures and missing data. This analysis applies multiple covariance structures to the data set and the structure with the lowest Akaike information criterion (AIC) was selected as best fit (Duricki et al., 2016; Verbeke, 1997). When two sessions were directly compared a two-way ANOVA or t-test was performed, as described below. For all analyses, statistical significance was set at *p* < 0.05, and Bonferroni *post hoc* comparisons were made when significant main effects or interactions were detected. Effect size (Cohen’s *d,* Cohen, 1988) was calculated for pairwise comparisons. Effect sizes were considered with respect to the following indices: 0.2, small; between 0.5 – 0.8, medium; between 1.2 – 2.0, large (Cohen, 1988; Sawilowsky, 2009).

LMM was conducted to compare experimental groups across sessions 1-3 (“laser on”) or sessions 4-6 (“laser off”). That is, sessions 1-3 or sessions 4-6 were used as the repeated variable and experimental group (control vs. halorhodopsin) was used as the between-subjects variable. For lever-directed behaviors, a LMM was also conducted to compare control sessions 1-3 (“laser on”) to halorhodopsin sessions 4-6 (“laser off”). A two-way ANOVA was conducted when session 3 (“laser on”) was directly compared to session 6 (“laser off”), with session (3 and 6) as the within subject independent variable and experimental group (control or halorhodopsin) as the between subject independent variable. Differences in the number of lever-cue contacts (dependent variable) between sessions 1 and 4 were analyzed using an unpaired t-test (control session 1 vs halorhodopsin session 4) or a paired t-test (control session 1 vs control session 4 or halorhodopsin session 1 vs halorhodopsin session 4). For both the LMM and ANOVA, lever-directed behaviors (number of lever-cue contacts, probability to approach the lever-cue, latency to approach the lever-cue), food magazine-directed behaviors (food magazine contacts during lever-cue presentation, probability to approach the food magazine during lever-cue presentation, latency to approach the food magazine during lever-cue presentation), and food magazine entries during the intertrial interval (ITI, non-CS food magazine head entries) were used as dependent variables.

Behavioral output from video analyses were also statistically analyzed. A two-way ANOVA was conducted to assess orienting responses directed towards the lever-cue or food magazine on session 3, with experimental group (control or halorhodopsin) and location (lever-cue or food magazine) as the independent variables. For the data generated by DLC, a Kolmogorov-Smirnov (KS) two-sample test compared the distributions for head direction responses (dependent variable) between experimental groups (control or halorhodopsin) at the following 4-s periods: 1) immediately before lever-cue presentation, 2) the first 4 s of lever-cue presentation, 3) the last 4 s of lever-cue presentation), 4) immediately after pellet delivery, and 5) during the inter-trial interval. Differences in approach bouts (each bout ≥ 1 s, dependent variable) between experimental groups (control or halorhodopsin) towards the lever-cue or food magazine 1) during lever-cue presentation and 2) after lever-cue retraction were analyzed using unpaired t-tests. Differences in locomotor activity and latency to approach the food magazine between experimental groups (control or halorhodopsin) were analyzed using unpaired t-tests. A two-way ANOVA was conducted to compare distance from the lever-cue during two time points: 1) 2 s before lever-cue presentation, and 2) 2 s after lever-cue presentation, with time as the within subject independent variable and experimental group (control or halorhodopsin) as the between subject independent variable.

## Results

### Behavioral characterization of the Long Evans transgenic rat colony

#### PavCA distribution

The tendency to sign- or goal-track (without optogenetic manipulation) was assessed in wild type (WT) and TH-Cre littermates from our in-house breeding colony. There were no significant differences in the PavCA Index across sessions 1-5 when comparing WT Naïve vs WT Sham vs TH-Cre (Table 1). Further, PavCA Index did not differ significantly across sessions between WT rats that did or did not receive sham surgery (Fisher’s exact test, *p* = 0.60); thus, these groups were collapsed for visualization (Figure 1b, c) and further analyses. While there were not robust differences between WT and TH-Cre rats across sessions, there was a significant group x session interaction (*F*_(4,172.373)_ = 3.175, *p* = 0.015) and post-hoc analyses revealed that TH-Cre rats had a higher PavCA Index on session 5 relative to WT rats (*F*_(1,93)_ = 4.328, *p* = 0.040, Cohen’s *d* = 0.46, Figure 1b, Table 1). Out of the total population of rats that were screened (N = 95), ∼84% were sign-trackers. Of the WT (n = 57 in total) rats ∼81% were sign-trackers and of the TH-Cre (n = 38 in total) rats, ∼89% were sign-trackers; but the population distribution between WT and TH-Cre rats was not significantly different (Fisher’s exact test, *p* = 0.40, Figure 1c). These data suggest that this colony of Long Evans male rats are skewed towards sign-trackers, regardless of genotype. This skew in the population provides an opportunity to assess the impact of neuronal manipulations on the attribution of incentive salience to reward cues and thereby the development of sign-tracking behavior.

**Table 1.** Statistical analyses for PavCA Index across sessions 1-5. Data from linear mixed effects model analyses for PavCA Index on sessions 1-5. The left panel compares WT Naïve, WT Sham, and TH-Cre animals. The right panel compares WTs (WT Naïve and WT Sham collapsed) to TH-Cre animals. Significant effects and interactions are bolded (\**p* < .05 and \*\*\**p* < .001).

|  | Sessions 1-5: PavCA Index |  |  |  |  |  |
| --- | --- | --- | --- | --- | --- | --- |
|  | WT Naïve vs WT Sham vs TH-Cre |  |  | WT vs TH-Cre |  |  |
|  | DF | F | p | DF | F | p |
| *Group* | 1,94.710 | 0.534 | 0.588 | 1,94.469 | 0.642 | 0.425 |
| *Session* | 1,155.351 | 51.978 | **\*\*\*<0.001** | 1,172.373 | 60.045 | **\*\*\*<0.001** |
| *Group\*Session* | 1,172.333 | 1.725 | 0.096 | 1,172.373 | 3.175 | **\*0.015** |

### Optogenetic inhibition of the VTA

#### Effects of optogenetic inhibition during lever-cue presentation in PavCA

To assess the role of dopamine in the attribution of incentive salience to a reward cue, we expressed an inhibitory opsin (eNpHR) in dopamine neurons in the VTA of TH-Cre rats (Figures 2 and 3). Disrupting cue-elicited dopamine through optogenetic inhibition of the VTA reduced lever-directed behaviors. During sessions 1-3, when the laser was turned on concurrently with lever-cue presentation, there was a significant effect of experimental group and/or a group x session interaction for all measures of lever-directed behavior (Table 2a). As shown in Figure 4a, while rats in the control group increased the number of lever-cue contacts across sessions 1-3, rats in the halorhodopsin group did not (group x session interaction: *F*_(2,15.773)_ = 4.119, *p* = 0.036; effect of session for control group: *F*_(2,15.279)_ = 4.418, *p* = 0.031). *Post hoc* comparisons revealed a significant reduction in the number of lever-cue contacts among halorhodopsin rats compared to control rats on session 3 (*p* = 0.02, Cohen’s *d* = 1.27) (Figure 4a). In agreement with this, the probability to approach the lever-cue was lower among halorhodopsin rats relative to control rats across sessions 1-3 (effect of group: *F*_(1,15.405)_ = 4.954, *p* = 0.041) and only those in the control group showed a significant increase in the probability to approach the lever-cue across sessions (group x session interaction: *F*_(2,18.848)_ = 5.237, *p* = 0.016; effect of session for control group *F*_(2,24.073)_ = 5.730, *p* = 0.009, Figure 4b). *Post hoc* analyses revealed that control rats had a significant increase in the probability to approach the lever-cue on session 3 relative to session 1 (*p* = 0.008, Cohen’s *d* = 0.95). Consistent with the results described above, the latency to approach the lever-cue significantly decreased across sessions 1-3 in control rats, but not those in the halorhodopsin group (effect of group: *F*_(1,15.396)_ = 4.620, *p* = 0.048, group x session interaction: *F*_(2,15.436)_ = 4.513, *p* = 0.029, effect of session for control group: *F*_(2,18.848)_ = 6.053, *p* = 0.009, Figure 4c). Again, *post hoc* comparisons revealed that the group differences were most apparent on session 3, when halorhodopsin rats took more time to approach the lever-cue relative to control rats (*p* = 0.008, Cohen’s *d* = 1.34; Figure 4c). Thus, for all these measures, the difference between groups was most apparent on session 3, after control rats began to exhibit sign-tracking conditioned response.

**Figure 4.**
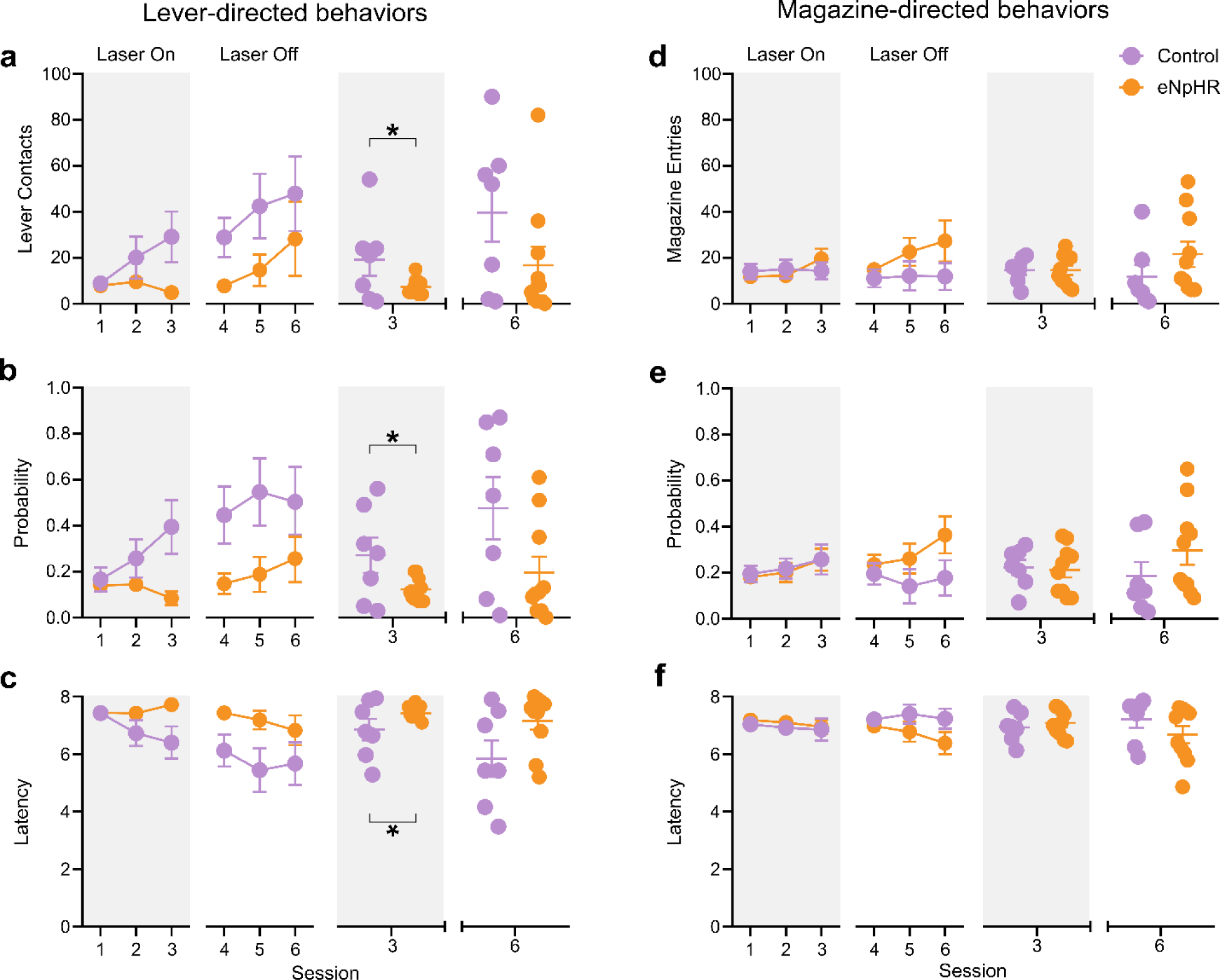
Inhibition of dopamine in the VTA attenuates sign-tracking behavior. (a-c) Lever-directed and (d-f) magazine-directed behaviors for sessions 1-3 (laser on) and 4-6 (laser off). (a-f) A comparison of session 3 (laser on) to session 6 (laser off) is to the right of each panel. Data are shown as mean ± SEM for a,d) number of contacts or head entries, (b,e) probability, or (c,f) latency to approach the lever-cue (left) or food magazine (right). (a-c) Optogenetic inhibition of dopamine neurons in the VTA decreases lever-directed behaviors in halorhodopsin rats (n = 10) compared to controls (n = 7) on sessions 1-3. Both groups had similar responding for lever-directed behaviors between sessions 4-6. (d-f) Optogenetic inhibition during sessions 1-3 had no effect on magazine-directed behaviors. Both groups had similar responding for magazine-directed behaviors between sessions 4-6. Bracket indicates significant difference between groups on session 3, *p<0.05.

**Table 2.**
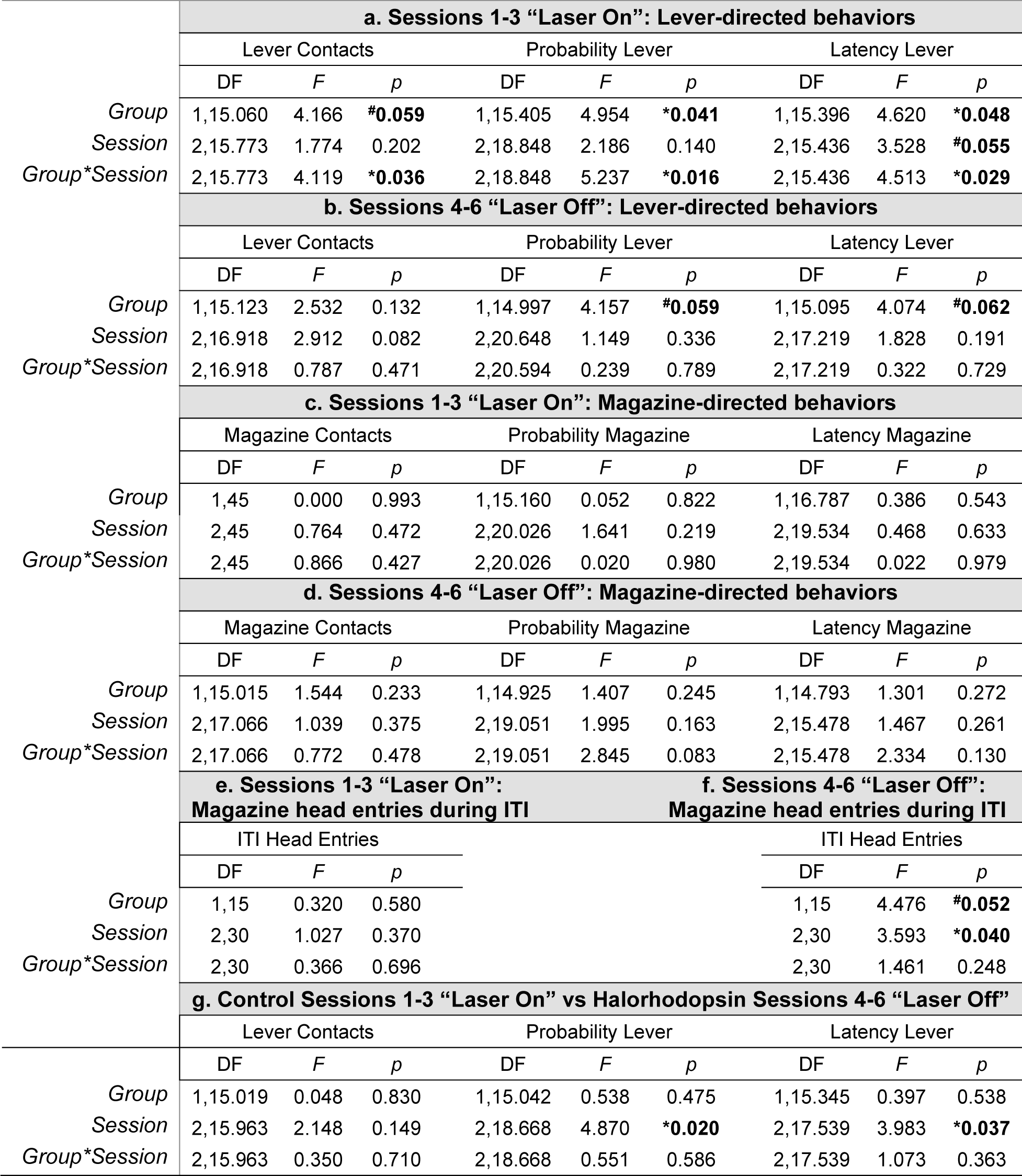
Statistical analyses for lever- and magazine-directed behaviors. Data from linear mixed effects model analyses for “laser on” and “laser off” sessions. Contacts, probability, and latency are represented for lever-directed behaviors, magazine-directed behaviors, and head entries into the magazine during the intertrial interval. Significant effects and interactions are bolded (^#^trend for a significant effect and \**p* < .05).

The impact of VTA dopamine inhibition was specific to lever-directed behaviors and did not affect goal-directed behaviors during sessions 1-3. Figures 4d-f illustrate the lack of differences between experimental groups for the number of food magazine contacts during lever-cue presentation, the probability to approach the food magazine during lever-cue presentation, and the latency to approach the food magazine during lever-cue presentation during “laser on” sessions (see also Table 2c).

Consistent with the data above, experimenter observation of the videos from session 3 revealed that the probability to orient towards the lever-cue significantly differed between experimental groups, whereas the probability to orient towards the food magazine did not (group x location interaction: *F*_(1,28)_ = 4.788, *p* = 0.039, effect of group for lever-cue: *F*_(1,24)_ = 5.538, *p* = 0.027; Figure 5a). As shown in Figure 5b, rats in the control group oriented to both the lever-cue and food-magazine on approximately 46% of trials, whereas those in the halorhodopsin group did so on approximately 36% of trials. Orientation to both the lever-cue and food cup on a given trial suggests that neither rats in the control group nor the halorhodopsin group were extreme sign-trackers or goal-trackers by session 3, and this is consistent with the data shown in Figure 4b. These data might also suggest that the value of the lever-cue has not been fully learned by session 3. Nonetheless, a conditioned orienting response is apparent for rats in both groups and inhibition of VTA dopamine selectively affects the response directed towards the lever-cue.

**Figure 5.**
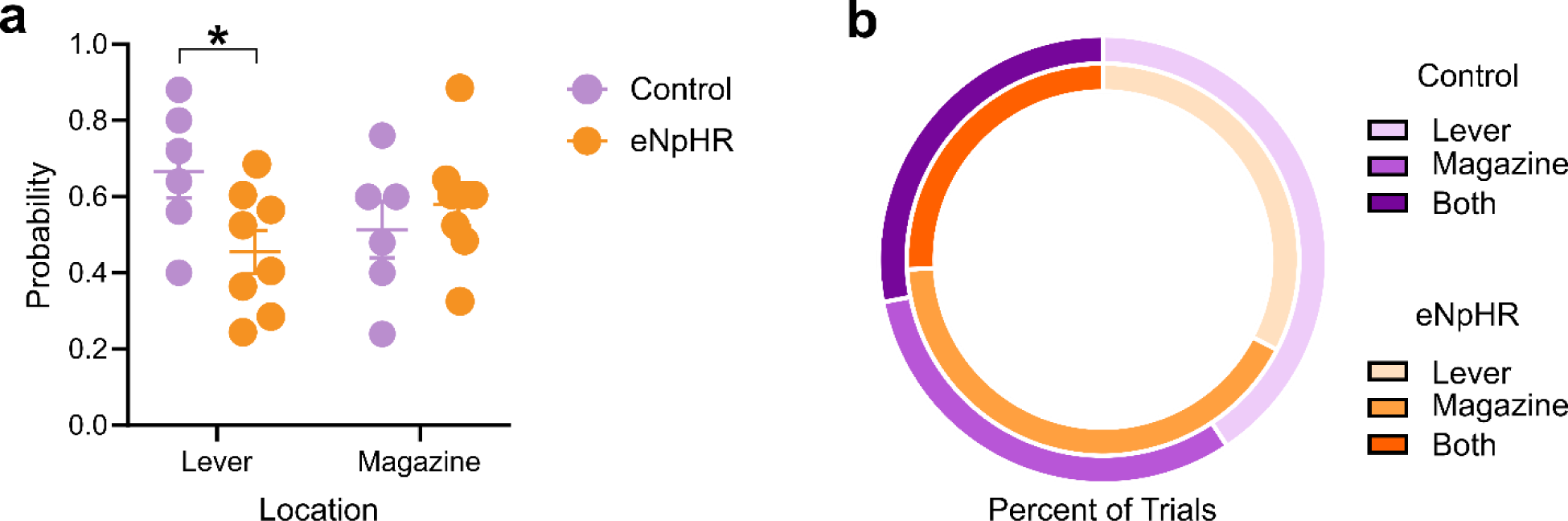
Head orientation to the lever-cue and food magazine during lever-cue presentation. (a) Mean ± SEM for probability to orient to the lever-cue or food magazine during lever-cue presentation on session 3 (final day of VTA dopamine inhibition). Rats in the control group oriented more towards the lever-cue than those that received VTA dopamine inhibition (i.e., eNpHR group). (b) Percent of trials to orient towards the lever-cue, food magazine, or both for control and eNpHR rats. Bracket indicates significant difference between groups on session 3, *p<0.05.

Importantly, the effects of VTA dopamine inhibition did not extend to behavior during the intertrial interval, as there were no significant differences in head entries into the food magazine in between trials during sessions 1-3 (Table 2e). Further, every rat consumed all of the food pellets that were delivered each session.

#### Effects of lifting optogenetic inhibition in later sessions

In the absence of VTA dopamine inhibition and laser presentation (sessions 4-6), there were no significant differences in lever-directed behaviors between halorhodopsin and control rats. For lever-directed behaviors, there were no significant differences between experimental groups for lever-cue contacts, probability to approach the lever-cue or latency to approach the lever-cue on sessions 4-6 (Figures 4a-c, Table 2b). Relative to rats in the halorhodopsin group, however, control rats showed a trend towards a greater probability to approach the lever-cue (effect of group: *F*_(1,14.997)_ = 4.157, *p* = 0.059) and a decreased latency to approach the lever-cue (effect of group: *F*_(1,15.095)_ = 4.074, *p* = 0.062; Figure 4b,c). As shown in Figure 4a-c, without VTA dopamine inhibition during lever-cue presentation, the halorhodopsin rats began to exhibit lever-directed behaviors comparable to control rats. In support, the “learning curve” for lever-directed behaviors did not differ between rats in the control group on sessions 1-3 and those in the halorhodopsin group on sessions 4-6 (Table 2g), when there was no laser inhibition. Further, there were no significant differences in lever-directed behaviors on session 1 for control rats relative to session 4 for halorhodopsin rats, and only rats in the control group had a significant increase in lever-cue contacts between session 1 and 4 (*t*_(6)_ = −2.62, *p* = 0.04). Taken together, these data demonstrate that rats in the halorhodopsin group did not attribute incentive salience to the lever-cue when they were receiving cue-paired laser inhibition, but once laser inhibition was removed, they were fully capable of doing so. Thus, VTA dopamine is necessary for encoding the incentive value of reward cues.

There were no significant group differences on sessions 4-6 in head entries, probability, or latency to approach the food magazine during lever-cue presentation (Figure 4d-f, Table 2d). While all rats tended to decrease the number of head entries into the food magazine during the intertrial interval across sessions 4-6 (effect of session: *F*_(2,30)_ = 3.593, *p* = 0.04, Table 2f) those in the control group tended to do so less than those with prior VTA dopamine inhibition (effect of group, *F_(_*_1,15)_ = 4.476, *p* = 0.05, Table 2f). These data suggest that rats in the halorhodopsin group may be less efficient in retrieving food pellets, and this is consistent with delayed entry into the food magazine upon lever-cue retraction, as presented below (Figure 9a).

#### Comparing “laser on” vs. “laser off” sessions

To further assess differences in behavior as a function of VTA dopamine inhibition, we directly compared session 3, the last PavCA session with laser inhibition, to session 6, the last PavCA session without laser inhibition. Rats in both the control group and halorhodopsin group showed an increase in lever-directed behaviors on session 6 relative to session 3 (Figure 4a-c, Table 3). Relative to rats in the halorhodopsin group, however, rats in the control group had a higher probability to approach the lever-cue (effect of group, *F_(_*_1,15)_ = 4.623, *p* = 0.048, Figure 4b) and had a tendency to do so more quickly (effect of group, *F_(_*_1,15)_ = 3.917, *p* = 0.066). There were no significant differences in food magazine-directed behaviors between session 3 vs. session 6 (Figure 4d-f).

**Table 3.**
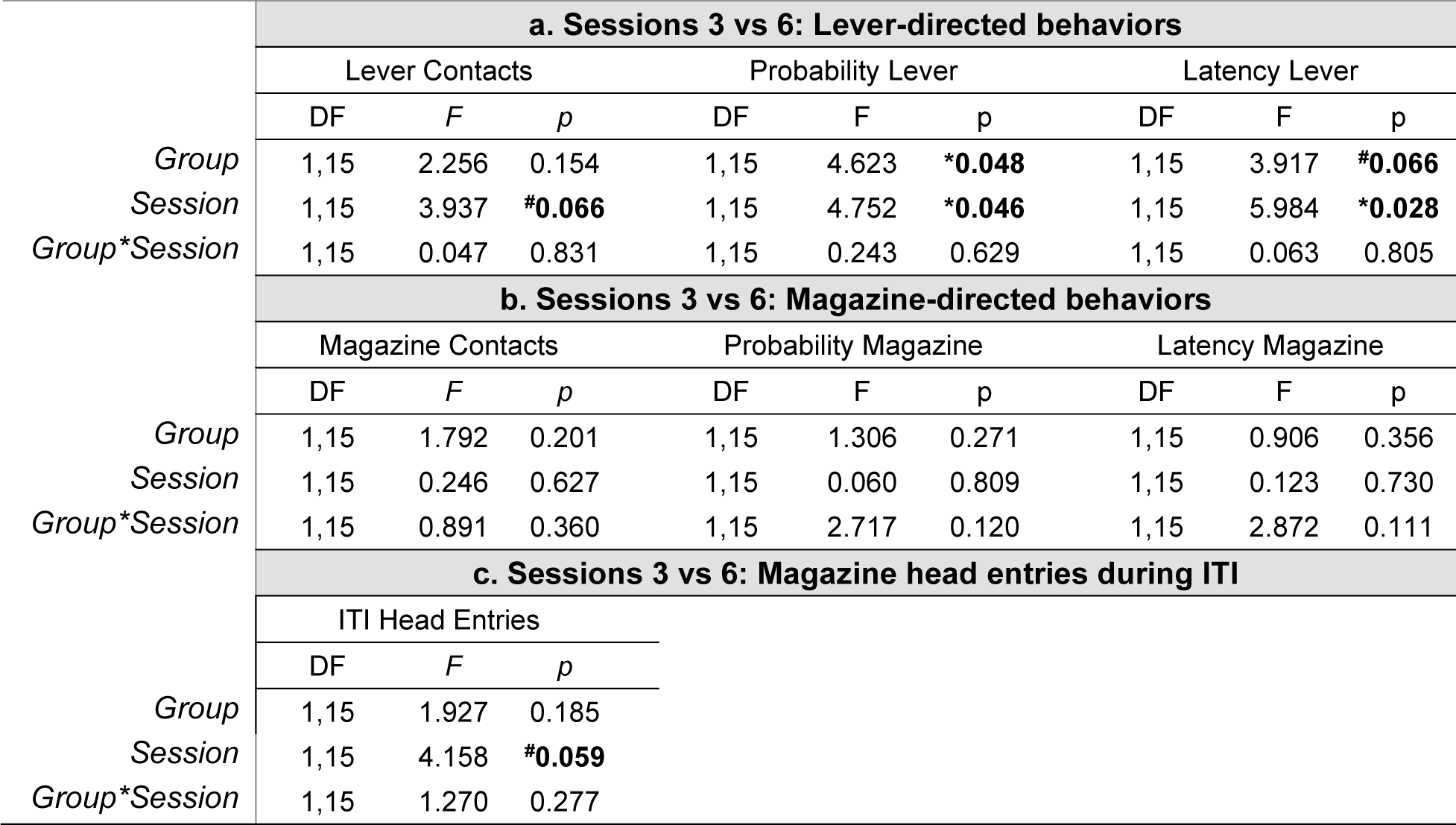
Statistical analyses comparing “laser on” vs “laser off” periods. Data from repeated measures ANOVA analyses comparing session 3 to session 6. Session 3 was the final session that animals received laser inhibition of dopamine neurons in the VTA and session 6 was the last session without laser inhibition. Statistical results for contacts, probability, and latency are represented for lever-directed behaviors and magazine-directed behaviors. Results for head entries during the intertrial interval are also shown. Significant effects are bolded (^#^trend for a significant effect, \**p* < .05, \*\**p* < .01, and \*\*\**p* < .001).

### Deep phenotyping expands behavioral analysis

Behavioral video analysis with DeepLabCut confirms that perturbing cue-elicited dopamine reduces multiple facets of cue-directed behaviors. Video analyses from session 3, the final session of VTA dopamine inhibition, were partitioned into different periods to assess location and time spent near (Figure 6), orientation to (Figure 7), and approach towards (Figure 8) the lever-cue or food magazine, as well as locomotor activity throughout the session (Figure 9).

**Figure 6.**
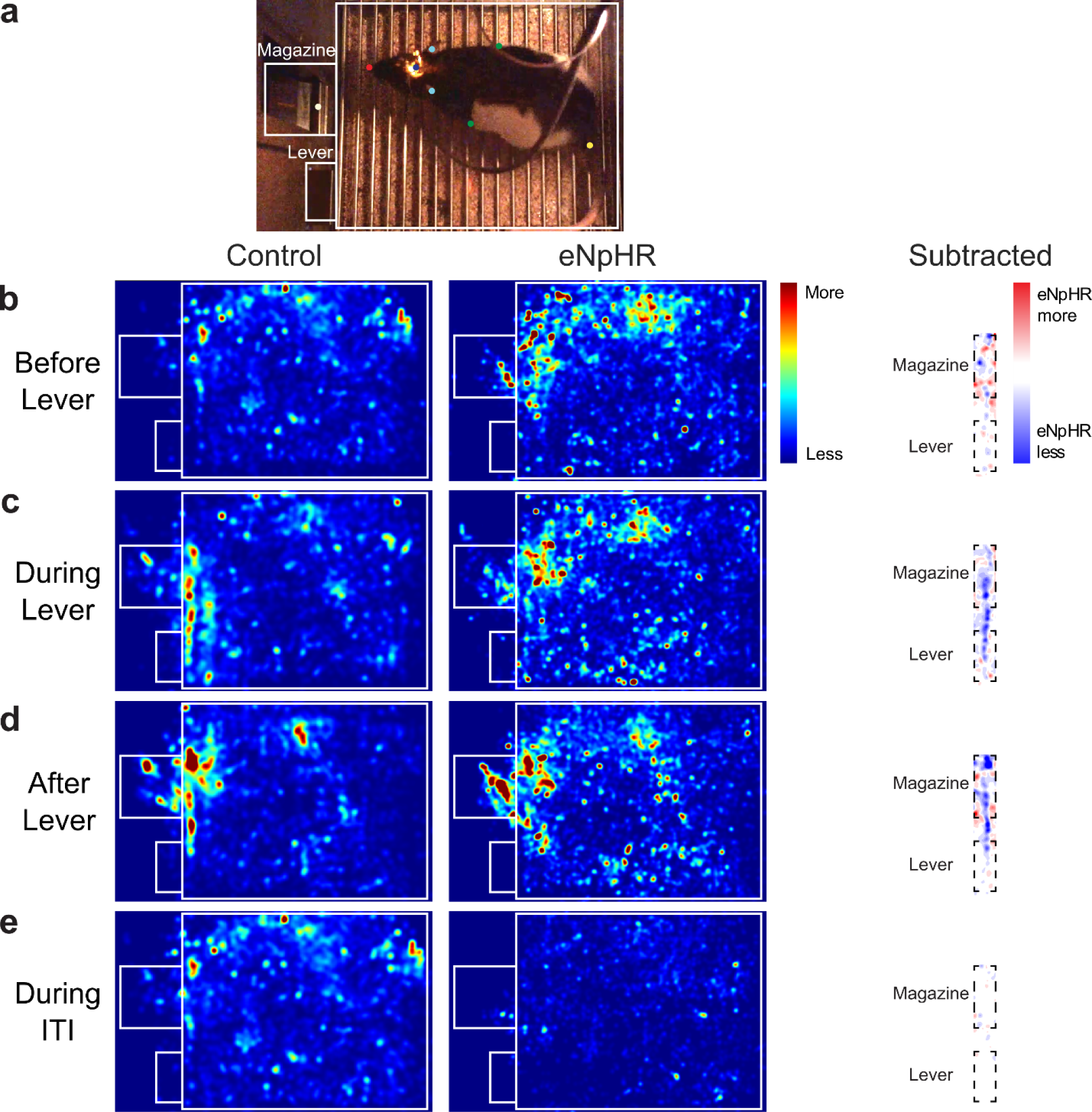
Inhibition of dopamine in the VTA reduces time spent near the lever-cue. (a) Representative frame from a video with arena borders overlayed and magazine/lever labeled. (b-e) Each heatmap represents the rats’ average location across all trials during session 3 of Pavlovian conditioned approach training (integrated over 123 video frames, 8.2-s time bins). Red indicates more time spent in a given location, blue indicates less. Location of the rats is shown for (b) the 8.2 s before lever-cue presentation, (c) 8.2 s during lever-cue presentation, (d) 8.2 s immediately after lever-cue retraction, and (e) 8.2 s during the inter-trial interval. Left: Control rats, Middle: eNpHR rats that received dopamine inhibition during lever-cue presentation, Right: Subtraction of eNpHR with control heat maps zoomed in on the magazine and lever. Red indicates that the eNpHR rats spent more time in a given location, blue indicates that control rats spent more time in a given location.

#### Location in chamber

Time spent in locations of the behavior chamber were analyzed in 8.2-s time bins, reflective of the period from lever-cue presentation to retraction. As indicated by the heatmaps shown in Figure 6b, in the 8.2 s period before the lever-cue was presented, control and halorhodopsin rats were found throughout the chamber with a tendency to gather near the food magazine. Once the lever-cue was presented rats in the control group appeared to spend more time near and around the lever-cue (Figure 6c), while those in the halorhodopsin group were around the food magazine or in other locations in the chamber. Immediately after lever retraction, rats in both groups spent more time at the location of food delivery (Figure 6d). A subtraction analysis further illustrates the difference in time spent near the lever-cue and food magazine for rats in the halorhodopsin group compared to those in the control group (Figure 6, right panel).

#### Head orientation

DeepLabCut analysis of head orientation revealed a trend toward a significant difference between groups in the 4-s preceding lever-cue presentation (KS test, *p* = 0.065), with rats in the control group showing a greater tendency to orient to the side of the chamber containing the lever-cue and food magazine (Figure 7b). During the first 4 s of lever-cue presentation, rats in the control group preferentially oriented towards the lever-cue relative to those in the halorhodopsin group (KS test, *p* = 0.034, Figure 7c). During the last 4 s of lever-cue presentation, there was a trend for a significant difference in head orientation between groups (KS test, *p* = 0.087, Figure 7c).There were no significant differences in head orientation to the lever-cue location or food magazine after pellet delivery (KS test, *p* = 0.328, Figure 7d) or during the intertrial interval (KS test, *p* = 0.118, Figure 7e). These data are consistent with those presented above and demonstrate that inhibition of VTA dopamine neurons impacts orientation towards the lever-cue upon its presentation, without affecting orientation towards the food magazine.

**Figure 7.**
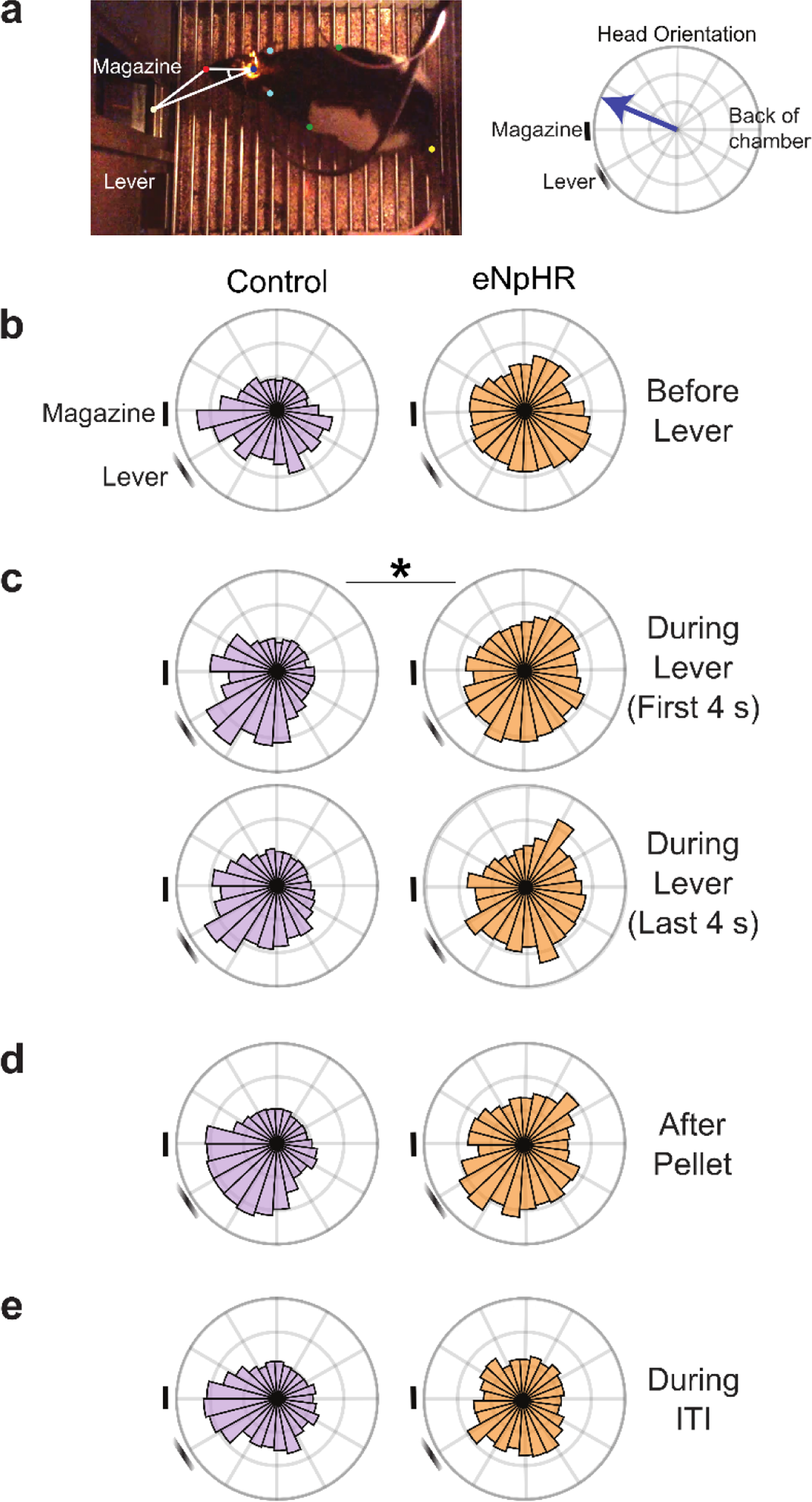
Inhibition of VTA dopamine neurons blunts the orienting responses to the lever-cue. (a) Left: Representative frame from a video with magazine and lever labeled. Marker labeling from Deep Lab Cut: Nose (red), tether (dark blue), ears (light blue), shoulders (green), tail (yellow), and magazine (white). Head orientation was calculated from the angle between two vectors: 1) from the tether to the nose and 2) from the tether to the magazine. Right: Head orientation was calculated from each video frame and is depicted as a unit vector on a polar histogram with the relative location of the magazine, lever, and back of chamber labeled. (b-d) Polar histograms showing average head direction (b) before lever-cue presentation (4 s before), (c) during lever-cue presentation (first and last 4 s), (d) after pellet delivery (4 s), and (e) during the inter-trial interval (ITI, 4 s). (c) Compared to controls, rats that received inhibition of dopamine neurons in the VTA had more variability in their head direction during lever-cue presentation. \**p* < 0.05.

#### Approach behavior

Approach towards the lever-cue and food magazine was assessed as an additional metric that is not captured by the Med Associates output, but one that is a hallmark of incentive salience attribution (Berridge, 2001; Cardinal et al., 2002). Approach behavior was counted when the rats’ nose was within 1 cm of either the lever-cue or food magazine for more than 1-s. Relative to rats in the control group, those in the halorhodopsin group showed less approach behavior towards the lever-cue upon its presentation (*t*_(12)_ = 4.911, *p* < 0.001, Figure 8a). Interestingly, the same was true during the intertrial intervals (i.e. after the lever had been retracted) (*t*_(12)_ = 3.753, *p* = 0.003, Figure 8b), reflecting a general tendency for rats in the control group to spend more time by the lever-cue location (see also Figure 6d). There were no significant differences between groups for approach towards the food magazine at any time point (i.e., during lever-cue presentation (*t*_(12)_ = −0.079, *p* = 0.939, Figure 8a) or after lever-cue retraction (*t*_(12)_ = −0.788, *p* = 0.446, Figure 8b).

**Figure 8.**
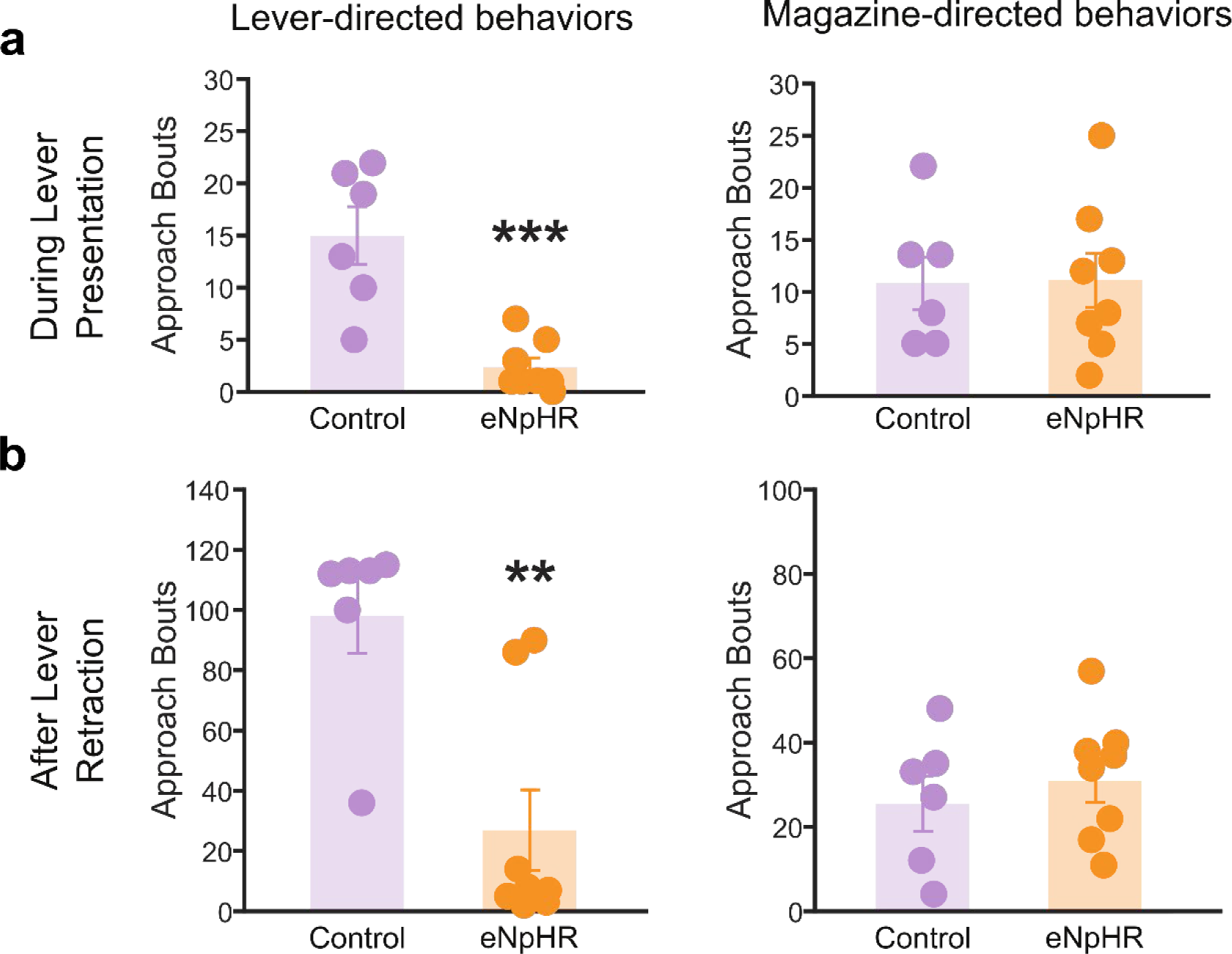
DeepLabCut analysis confirms inhibition of VTA dopamine neurons suppresses sign-tracking behavior. Data are shown as mean ± SEM for approach bouts. Inhibition of dopamine neurons reduces the amount of approach behavior towards the lever-cue, but not the food magazine, (a) during lever-cue presentation and (b) after lever-cue retraction (during the inter-trial intervals). \*\**p* < 0.01, and \*\*\**p* < 0.001.

#### Locomotor activity

Importantly, the experimental groups did not display significant differences in non-specific locomotion during session 3 (*t*_(12)_ = 0.406, *p* = 0.692, Figure 9a). When lever-cue-elicited locomotion was evaluated as the distance from the lever-cue 2 s before and 2 s after presentation there was a significant effect of time (*F*_(1,836)_ = 115.680, *p* < 0.001), but as shown in Figure 9b, the lever-cue-elicited locomotion was more apparent in rats in the control group relative to those in the halorhodopsin group (effect of group: *F*_(1,836)_ = 1135.491, *p* < 0.001, group x time interaction: *F*_(1,836)_ = 47.958, *p* < 0.001, Figure 9b). *Post hoc* comparisons revealed that rats in the control group moved closer to the lever-cue relative to those in the halorhodopsin group both before (*p* < 0.001, Cohen’s *d* = 0.01) and after lever-cue presentation (*p* < 0.001, Cohen’s *d* = 0.045). Interestingly, relative to those in the control group, rats in the halorhodopsin group were delayed in approaching the food magazine once the lever-cue had been retracted (*t*_(12)_ = −2.409, *p* = 0.03, Figure 9c). This delay could potentially be due to prolonged effects of VTA dopamine inhibition. Nonetheless, all rats consumed all of the food pellets and there were no significant differences between groups in general locomotor activity. Thus, these data support the notion that cue-elicited dopamine activity in the VTA plays a selective role in encoding the incentive value of reward cues.

**Figure 9.**
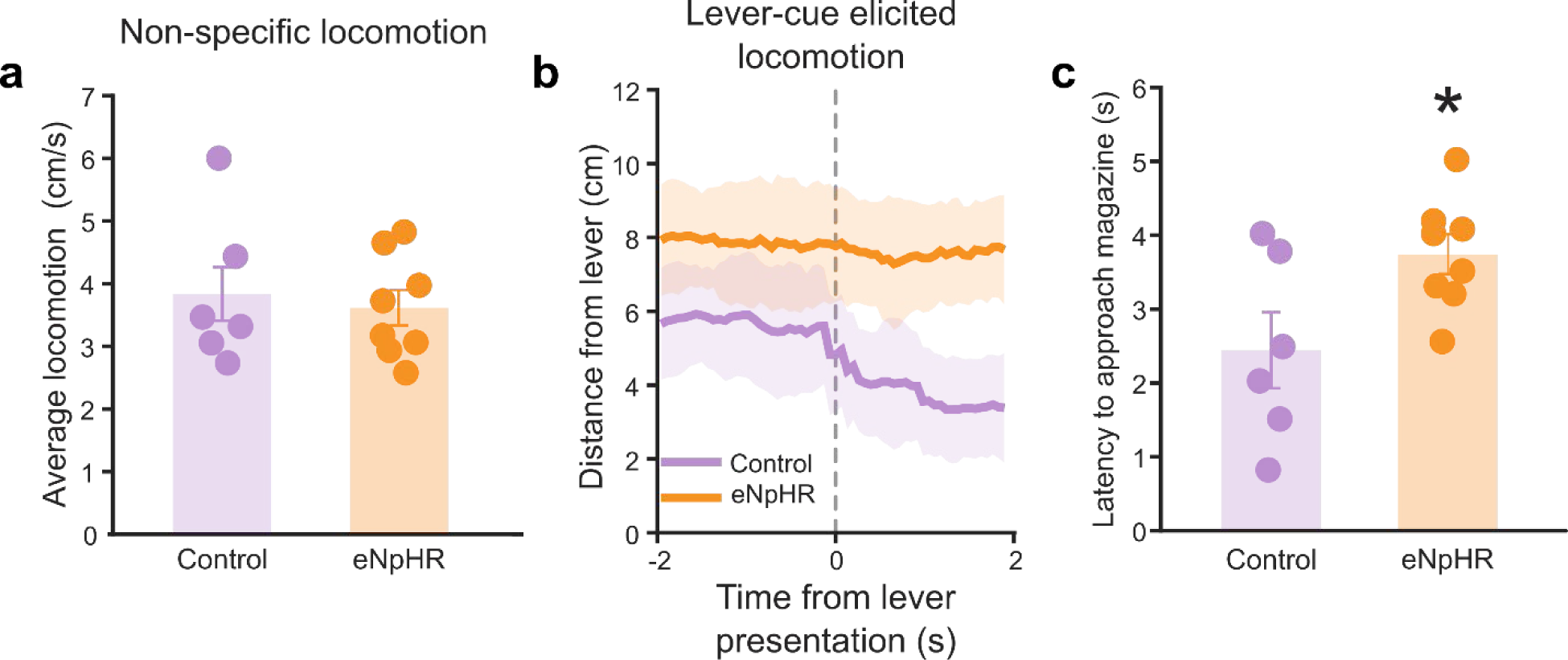
Inhibition of dopamine neurons does not change non-specific locomotion but does suppress cue-elicited locomotion. Data are shown as mean ± SEM. (a) Rats receiving inhibition of dopamine neurons showed no differences in non-specific locomotor activity as compared to controls. (b) Control rats moved closer to the lever-cue when it was presented relative to those that received inhibition of dopamine neurons. (c) Rats receiving inhibition of dopamine neurons were delayed in approaching the food magazine. \**p* < 0.05

## Discussion

It is known that VTA dopamine is involved in reward processing, however, the precise contributions of dopamine in terms of temporal specificity and value encoding remain a subject of debate (for review see: Berke, 2018; Berridge, 2007; Berridge, 2012; Stauffer, 2018; Triche et al., 2022; Zhang et al., 2009). Here we capitalized on the temporal precision of optogenetics and utilized a transgenic rat colony with a tendency to sign-track to further explore the role of dopamine in reward learning. We demonstrate that inhibition of VTA dopamine activity during presentation of a discrete cue that predicts reward delivery prevents incentive value encoding. Specifically, inhibition of VTA dopamine during lever-cue presentation precludes the attribution of incentive motivational value to the reward cue and thereby blocks the development of a sign-tracking conditioned response. Detailed analysis of behavior using DeepLabCut reinforced the specificity of these effects, revealing that locomotor activity was not affected by selective inhibition of VTA dopamine activity, nor was orientation or approach directed towards the location of reward delivery. Further, when VTA dopamine activity was restored, rats developed a sign-tracking conditioned response indicative of incentive salience attribution. These data are in agreement with prior studies demonstrating that dopamine is essential for incentive learning and the acquisition and expression of sign-tracking behavior (Chow et al., 2016; Flagel et al., 2011; Saunders & Robinson, 2012; Yager et al., 2015).

The role of dopamine in reward processing has been presented within the context of multiple learning theories and frameworks, some of which are in direct opposition (e.g., Berridge, 2007; Berridge, 2012; Gershman & Uchida, 2019; Langdon et al., 2018; Lerner et al., 2021; Schultz, 2019). The long-prevailing view that dopamine encodes the predictive value of reward cues and reflects a universal learning signal (Schultz et al., 1997) has been met with conflicting data. With the use of new technologies to further probe the role of dopamine in reward learning (for review see de Jong et al., 2022), it has been elegantly demonstrated that dopamine promotes associative learning (Sharpe et al., 2020) and encodes perceived saliency independent of valence (Kutlu et al., 2021), even when conditions are ripe for prediction error signals. Further, using a novel computational approach, it was shown that dopamine conveys causal associations without reward prediction error (Jeong et al., 2022). Our prior work with the sign-tracker/goal-tracker animal model is consistent with these more recent findings. We demonstrated that dopamine in the core of the nucleus accumbens encodes the incentive properties of reward cues, not the predictive (Flagel et al., 2011). That is, the classic prediction error shift in dopamine from the reward (unconditioned stimulus) to the reward cue (conditioned stimulus) occurs in sign-trackers, but not goal-trackers. If dopamine were merely a predictive learning signal, the shift in dopamine would have been evident in both sign-trackers and goal-trackers, as the reward-cue is a predictor and elicits a conditioned response for both. Moreover, blockade of dopamine signaling prevented the learning of a sign-tracking conditioned response, but not goal-tracking (Flagel et al., 2011; Saunders & Robinson, 2012).

Here we demonstrate that inhibition of VTA dopamine neurons selectively during lever-cue presentation prevents the attribution of incentive salience to the lever-cue and thereby precludes the development of a sign-tracking response. Upon restoration of VTA dopamine activity, the same rats attributed incentive value to the lever-cue in a manner that was indistinguishable from rats in the control group during their initial Pavlovian training sessions. Consistent with these findings, dopamine neurons in the VTA are activated to a much greater extent in sign-trackers relative to goal-trackers during lever-cue interaction (Ferguson et al., 2020) and VTA dopamine stimulation can itself transform a predictive stimulus into an incentive stimulus (Saunders et al., 2018). Specifically, neurons projecting from the VTA to the core of the nucleus accumbens were found to be especially important for incentive value encoding (Saunders et al., 2018); but others have reported that dopamine within the nucleus accumbens shell encodes incentive salience (Saddoris et al., 2015). The current findings add to the growing body of literature that supports a role for dopamine neurons in the VTA in incentive value encoding.

As the rats used in this study have an inherent predisposition to sign-track, we were not able to directly assess the effects of VTA dopamine inhibition on the development of goal-tracking behavior. Thus, dissociating the effects of cue-paired inhibition of VTA dopamine neurons on encoding the predictive versus incentive value of reward cues is complex. To better assess the effects of this manipulation on predictive learning we evaluated the conditioned orienting response, which is indicative that a stimulus-reward relationship has been learned (Buzsaki, 1982). Experimenter observation and DeepLabCut analyses revealed that rats in both groups exhibited a conditioned orienting response directed towards the lever-cue and/or food magazine, but only lever-cue oriented responses were affected by VTA dopamine inhibition. These findings are seemingly in contrast to prior studies that have shown that the conditioned orienting response directed towards the lever-cue remains intact in sign-trackers following administration of a dopamine antagonist into the core of the nucleus accumbens, and that only approach and interaction with the lever-cue is attenuated (Saunders & Robinson, 2012; Yager et al., 2015). It is important to note, however, that the manner in which we assessed conditioned orienting differs from these studies. For example, rats in the current study were not habituated to the presentation of the lever-cue in the absence of reward delivery (as in Yager et al., 2015).

Further, we assessed conditioned orienting behavior across all trials on sessions 3, presumably as the value of the lever-cue was still being learned, whereas other studies assessed it at the time an extreme sign-tracking response was evident (Saunders & Robinson, 2012; Yager et al., 2015) and only during the latter half of the session (Saunders & Robinson, 2012). It is also possible that, in the current study, the lever-directed conditioned orienting response is affected via inhibition of dopamine activity in non-striatal regions, such as the prefrontal cortex (Swanson, 1982), which is known to play a role in “cognitive” or model-based learning (Dayan & Balleine, 2002; Dickinson & Balleine, 2002; Ioanas et al., 2022; Kuhn et al., 2018; Morrens et al., 2020). Regardless, we believe our findings are consistent with the conclusion that dopamine is necessary for attributing incentive salience and transforming a predictive stimulus into a “motivational magnet”, but not for learning the stimulus-reward relationship. Based on these findings and those from other groups (Flagel et al., 2011; Saunders & Robinson, 2012), we would not expect cue-paired inhibition of VTA dopamine neurons to affect the prepotent conditioned response in goal-trackers.

Although locomotor activity was not affected by VTA dopamine inhibition in the current study, it is possible that this manipulation affected general motivation in real time. Rats that received cue-paired inhibition showed little change in locomotor activity during inhibition and were then slower to approach the location of reward delivery upon cessation of inhibition. The delay in approaching the food magazine upon lever-cue retraction could be a result of prolonged effects of dopamine inhibition or, alternatively, suggest that the perceived meaning of lever-cue retraction was still being learned. In support of the latter, rats in the halorhodopsin group continued to show increased responding in the food magazine relative to those in the control group during the intertrial interval. Further, others have reported that, relative to goal-trackers, sign-trackers show greater engagement of VTA dopamine neurons upon both the presentation and retraction of the lever-cue (Ferguson et al., 2020). It is also possible that inhibition of VTA dopamine neurons attenuated any incentive motivational value placed upon the food magazine (DiFeliceantonio & Berridge, 2012; Mahler & Berridge, 2009). Importantly, however, the fact that a conditioned orienting response directed towards the food magazine did not differ between groups and that all of the rats consumed all of the food pellets that were delivered suggests that lever-cue-paired VTA dopamine inhibition does not generally affect learning or motivation.

The current findings expand and enhance the existing literature pertaining to the role of dopamine in reward learning. We clearly demonstrate that lever-cue-paired inhibition of VTA dopamine activity prevents the attribution of incentive motivational value to the reward cue and the development of sign-tracking behavior. Using DeepLabCut, we were able to thoroughly assess metrics of learning and incentive motivation and rule out non-specific effects of VTA dopamine inhibition. This study was designed around the fact that the TH-Cre population we used was skewed towards sign-trackers; yet, this precluded us from assessing the effects of this manipulation in goal-trackers. The findings were interpreted with this individual variability or lack thereof in mind, and we consider this a valuable example for the field to consider. Further, the complexity of behavior and nuances therein are illustrated here and should be noted for those using cutting-edge techniques to decipher brain-behavior relationships. We will continue to capitalize on “deep phenotyping” approaches to assess the effects of similar manipulations in female rats and to better elucidate the neural substrates underlying reward learning and incentive value encoding. Based on the current findings, however, we conclude that cue-elicited dopamine is critical for the attribution of incentive salience to reward cues.

## Acknowledgements

The work was supported by NIH R01 DA038599 (S.B.F.) and DA054094 (S.B.F.), the Hope for Depression Research Foundation (H.A.), a NARSAD Young Investigator Grant (C.R.B.), and NIH R01 DK129366 (C.R.B.). A.G.I. and A.S.C. were supported by a NIDA T32 Training Program in Neuroscience (NIH T32-DA7281), A.G.I. was also supported by a National Science Foundation Graduate Research Fellowship (DGE 1256260), and a Rackham Merit Fellowship, University of Michigan. Use of DeepLabCut was supported in part through computation resources and services provided by Advanced Research Computing at the University of Michigan, Ann Arbor. We would like to thank Drs. Kent Berridge, Stephen Chang, and Terry Robinson for their insightful comments on earlier versions of this manuscript.

